# Can *Cynodon dactylon* be used to suppress invasive weeds? The effects of density-dependent on the growth and development of *Tagetes minuta* and *Gutenbergia cordifolia*

**DOI:** 10.1101/674085

**Authors:** Issakwisa B. Ngondya, Anna C. Treydte, Patrick A. Ndakidemi, Linus K. Munishi

## Abstract

Plant-Plant competitive interactions have been reported to be among the forces that shape plant community structure. We studied the effects of varying the density of *Cynodon dactylon* on the growth and development of the invasive plant species *Tagetes minuta* and *Gutenbergia cordifolia* in pot and field plot experiments following a completely randomized design. Increasing densities of *C. dactylon* strongly reduced *T. minuta* and *G. cordifolia* growth and development, leaf total chlorophyll and increased leaf anthocyanin of both invasive species. These detrimental effects may have contributed to poorer *T. minuta* and *G. cordifolia* performance under *C. dactylon* densities of more than 8 individuals per pot/plot compared to those in pots/plots without *C. dactylon*. This study suggests that *C. dactylon* can be successfully used to manage the two invasive plants, thus, improving forage production and biomass in affected rangelands.

## 1.0. Introduction

Since the 19^th^ century, the mechanism by which plants influence the structure and composition of their surrounding plant community has been investigated. Competition has since then received a lot of attention in ecological research [1–5] and was found to directly affect the local distribution of plants in a community [6]. Plant-Plant competition has well been demonstrated in a range of ecosystems; most vividly in ecosystems where native plants have been exposed to several stresses, for instance water shortage, soil nutrient deprivation and ecological invasion [7]. The most competitive plant always dominates the ecosystem and hence, poses a risk for local extinction of some associated flora and fauna.

The high competition ability for nutrients, water and light of some native grass species such as *Cynodon dactylon* is likely to contribute to their competitiveness [8]. *Cynodon dactylon* has been reported to successfully escape from stresses like invasion and drought by creeping away from invaded areas through stolones and by developing a deep root system [9,10]. It can grow on soils with a wide range of pH, survive flooding [11] and can grow over twice as large in mixed cultures than in monoculture [12]. The plant has further been reported to be highly competitive over most crops [13], which highlights its importance as a potential fodder grass for management of invasive weeds.

*Tagetes minuta* is an unpalatable exotic invasive plant native to Mexico [14] and it has escaped cultivation in most nations and is considered a noxious weed in parts of southern Africa [15]. This species has been introduced into various areas to the extent that it became a weed in most rangelands and farmlands of Tanzania [16]. Recently, the species has been reported as among problematic weeds that have invaded the Ngorongoro Conservation Area [17]. *Gutenbergia cordifolia* on the other hand is an annual plant native to Africa. Its leaves and flowers are allergenic and toxic to animals as they contain a chemical sesquiterpene lactone [18,19] that alters the microbial composition of the rumen and thereby affects its overall metabolic functioning [20]. While in Kenya the plant has already been reported as an invasive weed in most farmlands [21,22] in Tanzania the species seems to have invaded and dominated more than half of the World Heritage site Ngorongoro crater floor (250 km^2^) [23] and most parts of the Serengeti ecosystem (Pers. obs).

Management of invasive weeds in protected ecosystems poses great challenges as herbicides, which have proven successful in farmlands, are not recommended. Herbicides often have strong negative effects on other native flora and fauna should they be chosen for management purposes [24,25]. Although alternative management options such as uprooting and mowing are often opted for, they are labor-demanding and only a short term remedy as many invasive plant seeds remain in the soil seed bank. Therefore, we claim that if utilized well, plant density dependent competitive interactions might present an opportunity for developing management strategies for some problematic weeds such as *Tagetes minuta* and *Gutenbergia cordifolia*, thereby helping in the restoration of previously invaded ecosystems particularly grazing areas. As a low-cost, low impact management technique, plant-plant competition has been reported to be effective in restoration projects [26].

We aimed at utilizing *C. dactylon* as a competitor due to its agronomic value as a forage species [27]. Also this species was found in previous studies to be highly competitive [8] due to, among other reasons, its ability to form deep roots [9,10] and as it can grow on soils with a wide range of pH [11]. We studied the density dependent competitive effect of *C. dactylon* on growth parameters and leaf pigments of *T. minuta* and *G. cordifolia*. We set up screen house and field plot experiments by varying *C. dactylon* densities and hypothesized that this species will suppress the two weeds and, therefore, reduce their growth and development through suppression of the studied parameters. This study will pave a way for the application of *C. dactylon* as a management strategy in controlling of *T. minuta* and *G. cordifolia* invaded areas in rangelands for improved pasture production.

## 2.0. Materials and Methods

### 2.1. Experimental design

*Tagetes minuta* and *G. cordifolia* seeds were sown separately in pots (screen house) and in plots (field), in combination with *C. dactylon* following a simple additive design [28] in early 2016. Clay-loam soil was used in both experiments, which is similar to the Ngorongoro Crater’s soil where invasion has occurred. The two invasive species were referred to as “Weed species (*W*)” i.e. *Wt* and *Wg* for *T. minuta* and *G. cordifolia*, respectively while *C. dactylon* was referred to as “species (*Cd*)”. Weed species were grown in combination with *Cd* in pots of 0.15 m height × 0.56 m diameter and plots of 0.50 m × 0.50 m (Fig 1).

**Fig 1.**
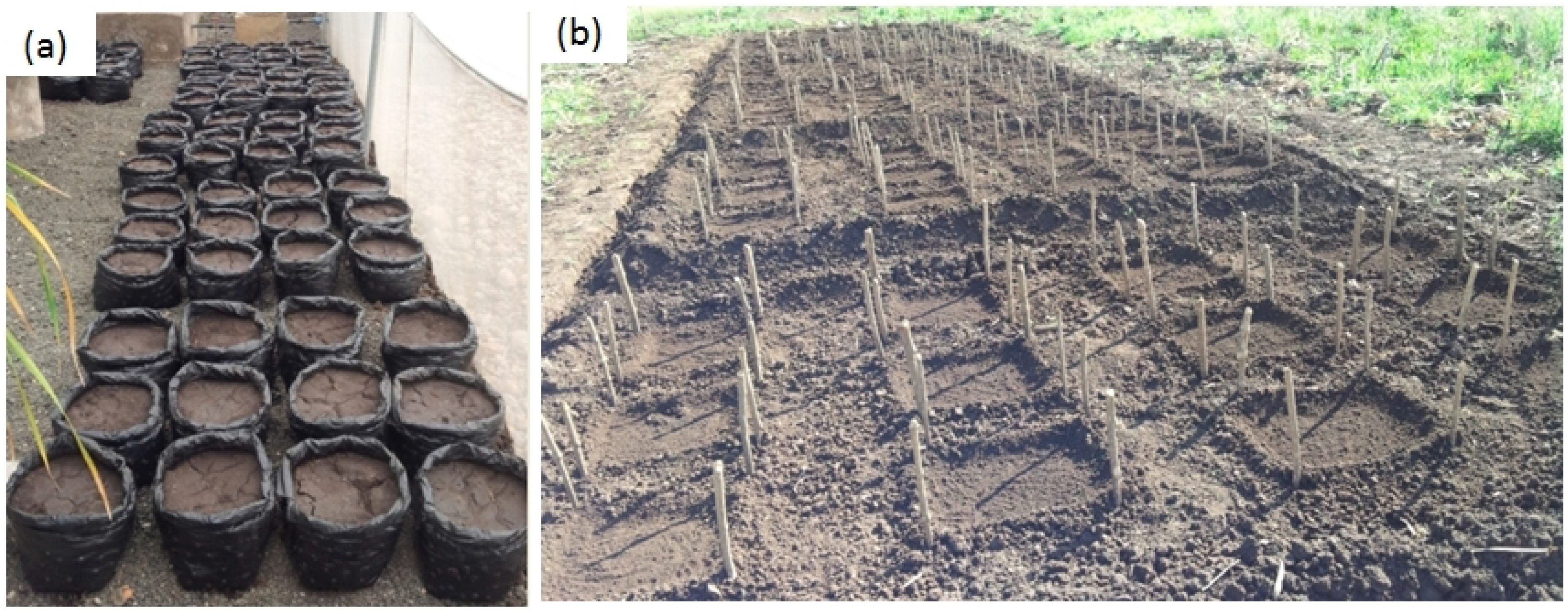
Pots (a) and plots (b) used in both screen house and field experiments respectively

Based on the used pots/plots size, density proportions of sown weed versus species Cd (W: Cd) were as follows: 2:0, 2:4, 2:6, 2:8 and 2:10, whereby 2:0 was used as control and each treatment was replicated three times (Fig 2). A total of 30 pots (five densities, three replications and two species) and 30 plots (five densities, three replications and two species) were used in this study. The interaction between *Wt*, *Wg* and *Cd* under uniform conditions (space, moisture and nutrients) was studied using a completely randomized design. Seeds of both *Cd* and *W* were sown at a spacing of ≥ 2 cm apart. During the first two weeks, 100% of planted seeds germinated, during this period pots / plots were irrigated with water ad-libitum to ensure establishment. After successful establishment plants in all pots / plots were irrigated with 0.5 liters of water daily, watering ceased at anthesis which marks the end of vegetative growth (meristematic and elongation) phase of both weeds (9 weeks).

**Fig 2.**
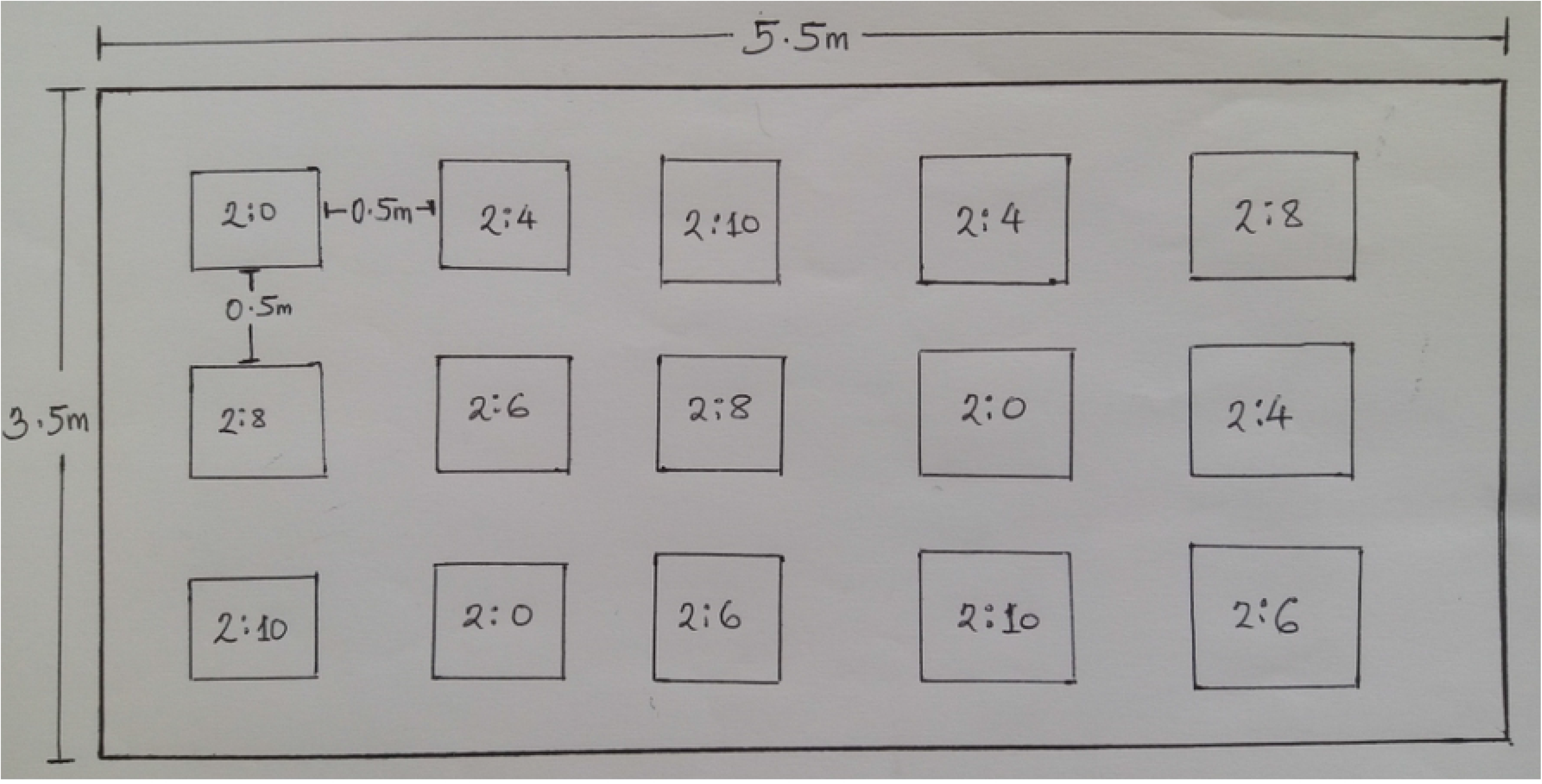
Experimental pot/ plots layout

### 2.2. Parameters measured

Number of vegetative branches, leaves, panicles, and seedling height and shoot diameter were measured parameters at the end of the plant’s vegetative growth phase. In this study, the number of weeds per pot/plot was considered as the entire population, 100% of which was sampled (two plants per pot/plot). The number of vegetative branches leaves and panicles were counted for each plant. Plant height was measured using a meter ruler while shoot diameter was measured using vernier calipers at a height of 5 cm from the ground. The total number of leaves in all pots / plots under the same treatment was considered as a population; over 30% of leaves were randomly sampled for leaf area determination. Leaf areas were determined using java based image processing software Image J (https://imagej.nih.gov/ij/) [29]. *Tagetes minuta* and *G. cordifolia* root lengths were measured using a meter ruler. Young leaves from the top-most part of the plant were sampled randomly per pot for chlorophyll determination while mature leaves were randomly sampled for anthocyanin level determination.

During the 9^th^ week, (the end of vegetative growth phase) both *T. minuta* and *G. cordifolia* were harvested (uprooted), washed, placed into paper bags and dried at 80°C for 48 hours [30]. Shoot and root material was separated and weighed to obtain total above / below ground dry biomass [30].

### 2.3. Measurement of leaf pigments

Leaf chlorophyll content has been linked to plant health status [31] and, therefore, a crucial parameter to be assessed during plant growth and development. Leaf chlorophyll of *T. minuta* and *G. cordifolia* plants was extracted according to [32] with some modifications: 50 mg of fresh leaves of 2.25 cm^2^ were immersed in 4 ml of Dimethyl Sulfoxide (DMSO) and incubated at 65°C for 12 h. The extract was transferred to glass cuvettes for absorbance determination. The absorbance of blank liquid (DMSO) and samples were determined under 2000 UV/VIS spectrophotometer (UNICO^®^) at 663 nm and 645 nm [32] and the total leaf chlorophyll (total Chl) calculated according to [33] using the following equation:

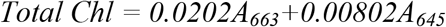

Where *A*_*663*_ and *A*_*645*_ are absorbance readings at 663 nm and 645 nm respectively

Bioassay of levels of anthocyanins in leaves of *T. minuta* and *G. cordifolia* were performed as described by [34]. Leaves of *T. minuta* and *G. cordifolia* were oven-dried at 60°C for 48 h, weighed, ground into a fine powder. Then, 0.10 g of leaf powder was weighed and mixed with 10 ml of acidified methanol prepared from a ratio of 79:20:1 MeOH:H_2_O:HCl. The mixture was incubated for 72 h in darkness for auto-extraction and filtered through Whatman paper Number 2. The extract was transferred to glass cuvettes for absorbance determination. The absorbance of acidified methanol as standard and that of samples were determined under a 2000 UV/VIS spectrophotometer (UNICO^®^) at 530 nm and 657 nm and expressed as Abs g.DM-1 [34]. Anthocyanin concentration in leaf extracts was measured as A_530_ - 1/3A_657_ [34] where *A*_*530*_ and *A*_*657*_ are absorbance readings at 530 nm and 657nm, respectively.

### 2.4. Data analysis

Shapiro-Wilk test for normality was performed on the number of vegetative branches, leaves, panicles, Plant height, shoot diameter, root length, leaf area, leaf total chlorophyll content, leaf anthocyanin concentration, shoot and root biomass of *T. minuta* and *G. cordifolia*. For all data that passed normality test, one-way analysis of variance (ANOVA) was carried out whilst for non-normally distributed data a Kruskal–Wallis test was performed. For both invasive weed species, one-way ANOVA was performed on the number of vegetative branches, number of panicles, leaf area, shoot diameter, leaf total chlorophyll and leaf anthocyanins concentration versus varying density of *C. dactylon*. Kruskal-Wallis test was carried out on the number of leaves, plant height, root biomass, shoot biomass and root length per plant. Pearson’s Product Moment and Spearman correlations were also performed on normally and non-normally distributed data respectively. The resulting means were separated by the Fisher’s Least Significant Difference (LSD). The statistical software used was STATISTICA version 8 and the level of significance was set at *p* < 0.05.

## 3.0. Results

### 3.1. Cynodon dactylon density dependent competitive effects on T. minuta and G. Cordifolia

#### 3.1.1. General observation

General decrease in *Gutenbergia cordifolia* and *Tagetes minuta* vigor was observed along increasing density gradient of *Cynodon dactylon* (Figs 3 and 4).

**Fig 3.**
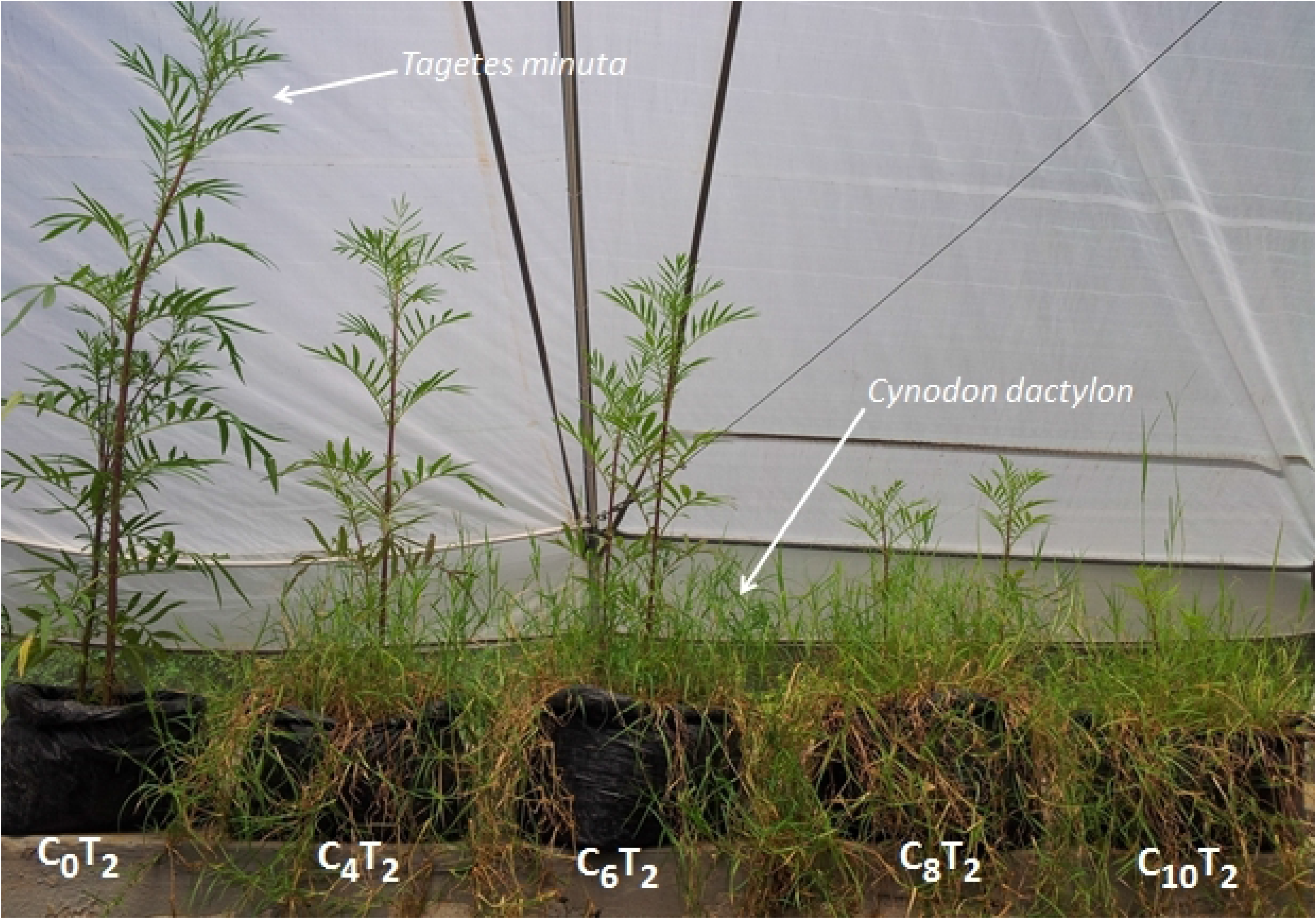
Effects of increasing densities of *Cynodon dactylon* on *T. minuta* vigor (C=*Cynodon dactylon* and T=*Tagetes minuta*, numbers represent proportions of sown *Cynodon dactylon* and *Tagetes minuta* where by C_0_T_2_ was a control)

**Fig 4.**
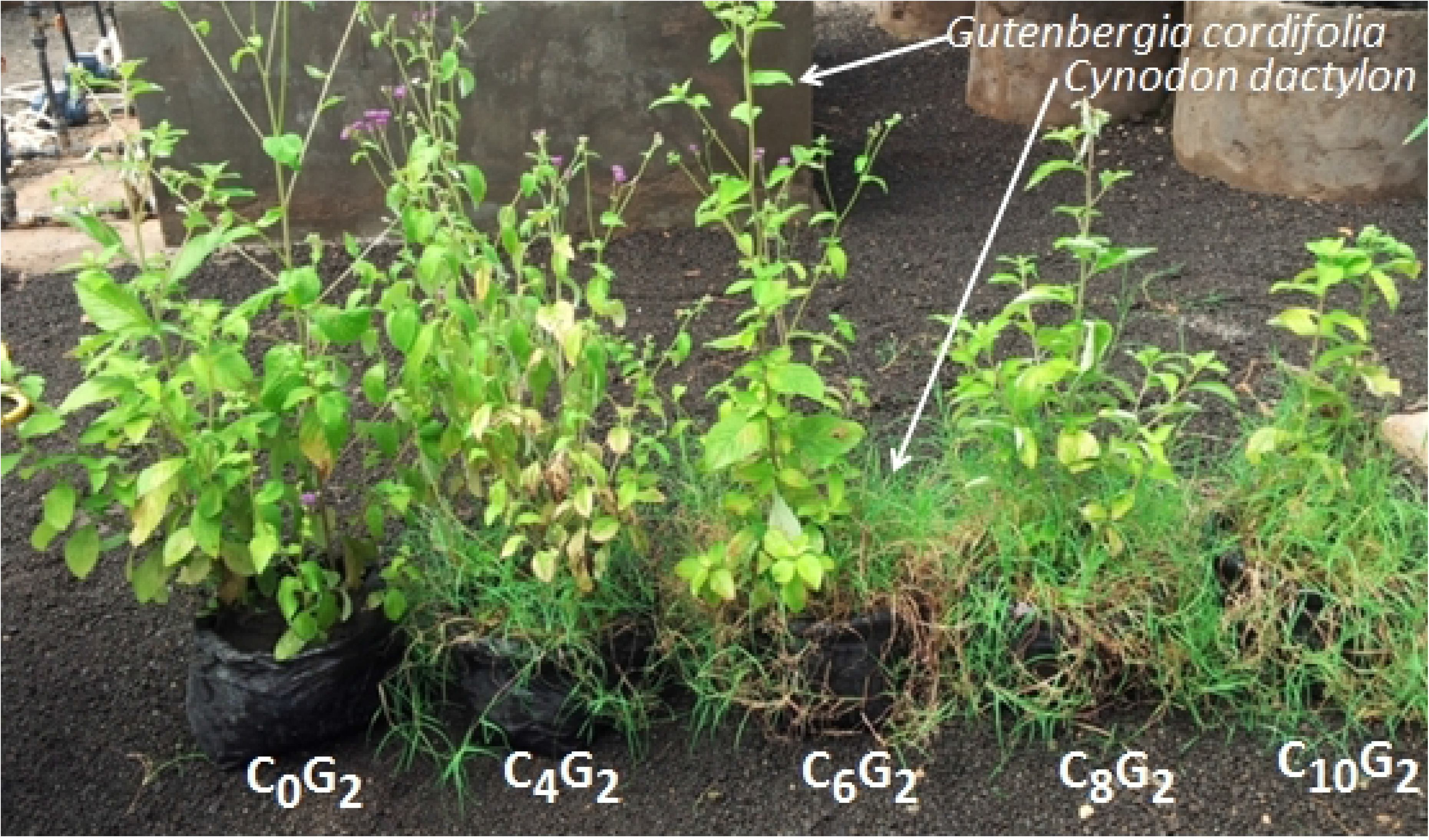
Effects of increasing densities of *Cynodon dactylon* on *G. cordifolia* vigor (C= *Cynodon dactylon* and G=*Gutenbergia cordifolia*, numbers represent proportions of sown *Cynodon dactylon* and *Tagetes minuta* where by C_0_G_2_ was a control)

#### 3.1.2. Plant growth parameters

The mean number of vegetative branches, panicles and leaf area of both *T. minuta* and *G. cordifolia* species differed significantly across the five *C. dactylon* treatments (Tables 1 and 2) and was over four times higher in control pots/plots than in pots/plots with *C. dactylon* when the latter’s densities reached more than 8 individuals per pot or plot (Figs 5 and 6).

**Table 1.**
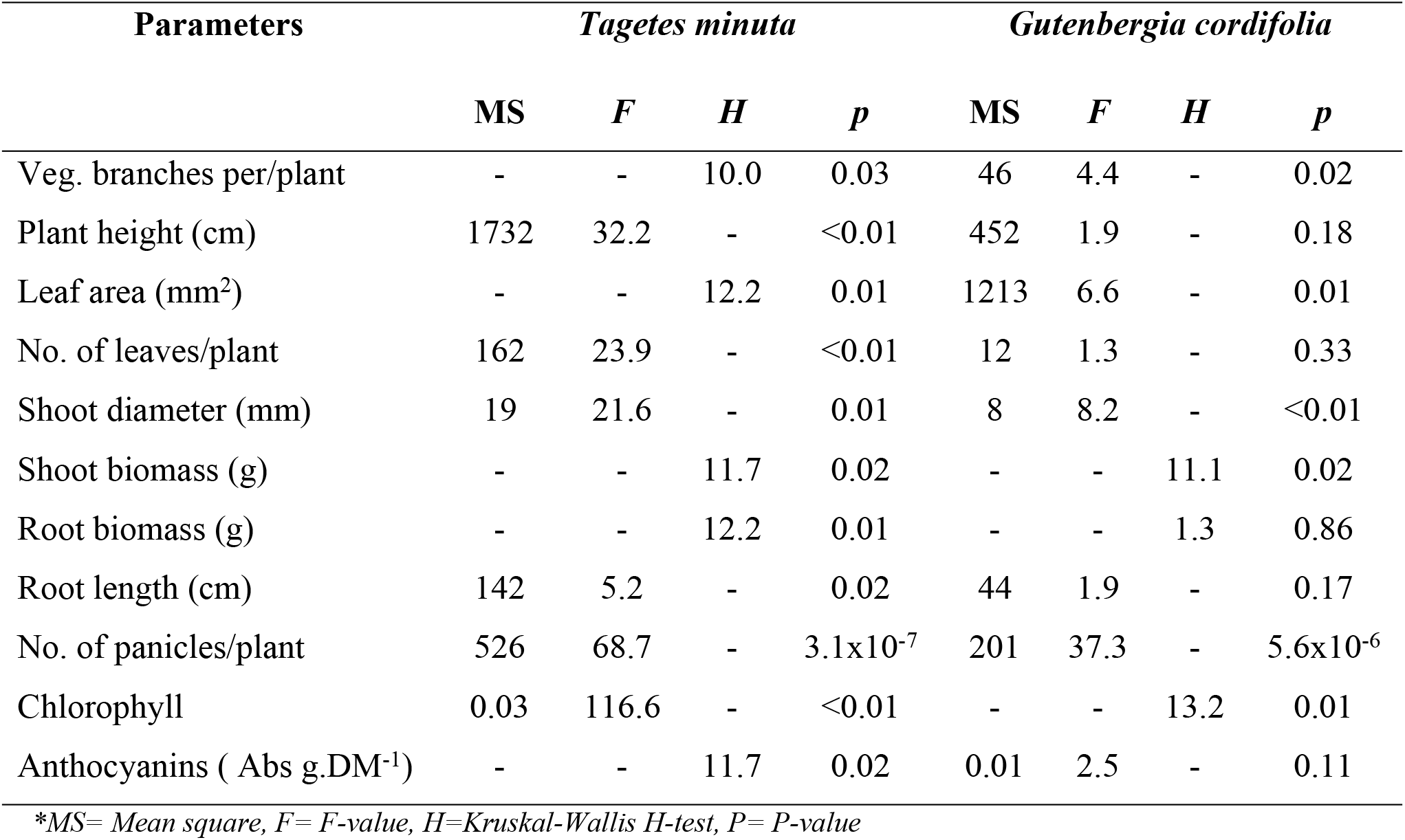
Effects of increasing densities of *Cynodon dactylon* on *T. minuta and G. cordifolia* growth parameters and leaf pigmentation in the screen house experiment

| Parameters | *Tagetes minuta* |  |  |  | *Gutenbergia cordifolia* |  |  |  |
| --- | --- | --- | --- | --- | --- | --- | --- | --- |
|  | MS | F | H | p | MS | F | H | p |
| Veg. branches per/plant | - | - | 10.0 | 0.03 | 46 | 4.4 | - | 0.02 |
| Plant height (cm) | 1732 | 32.2 | - | <0.01 | 452 | 1.9 | - | 0.18 |
| Leaf area (mm<sup>2</sup>) | - | - | 12.2 | 0.01 | 1213 | 6.6 | - | 0.01 |
| No. of leaves/plant | 162 | 23.9 | - | <0.01 | 12 | 1.3 | - | 0.33 |
| Shoot diameter (mm) | 19 | 21.6 | - | 0.01 | 8 | 8.2 | - | <0.01 |
| Shoot biomass (g) | - | - | 11.7 | 0.02 | - | - | 11.1 | 0.02 |
| Root biomass (g) | - | - | 12.2 | 0.01 | - | - | 1.3 | 0.86 |
| Root length (cm) | 142 | 5.2 | - | 0.02 | 44 | 1.9 | - | 0.17 |
| No. of panicles/plant | 526 | 68.7 | - | 3.1x10<sup>-7</sup> | 201 | 37.3 | - | 5.6x10<sup>-6</sup> |
| Chlorophyll | 0.03 | 116.6 | - | <0.01 | - | - | 13.2 | 0.01 |
| Anthocyanins ( Abs g.DM<sup>-1</sup>) | - | - | 11.7 | 0.02 | 0.01 | 2.5 | - | 0.11 |

\*MS= Mean square, F= F-value, H=Kruskal-Wallis H-test, P= P-value

**Table 2.** Effects of increasing densities of *Cynodon dactylon* on *T. minuta and G. cordifolia* growth parameters and leaf pigmentation in the field plot experiment.

| Parameters | *Tagetes minuta* |  |  |  | *Gutenbergia cordifolia* |  |  |  |
| --- | --- | --- | --- | --- | --- | --- | --- | --- |
|  | MS | F | H | P | MS | F | H | P |
| Veg. branches per/plant | 379 | 65.1 | - | <0.01 | 184 | 14.3 | - | <0.01 |
| Plant height (cm) | 1414 | 13.3 | - | 0.01 | 480 | 1.7 | - | 0.22 |
| Leaf area (mm<sup>2</sup>) | 43487857 | 31.4 | - | <0.01 | - | - | 12.3 | 0.02 |
| No. of leaves/plant | 206 | 17.4 | - | <0.01 | 67 | 7.9 | - | 0.01 |
| Shoot diameter (mm) | 35 | 38.7 | - | <0.01 | 9.7 | 32.8 | - | <0.01 |
| Shoot biomass (g) | - | - | 13.5 | 0.01 | - | - | 10.5 | 0.03 |
| Root biomass (g) | - | - | 13.2 | 0.01 | - | - | 7.0 | 0.13 |
| Shoot biomass (g) | - | - | 13.5 | 0.01 | - | - | 10.5 | 0.03 |
| Root length (cm) | 250 | 13.8 | - | <0.01 | 54.8 | 2.8 | - | 0.08 |
| No. of panicles/plant | 439 | 41.6 | - | 3.3x10<sup>-6</sup> | - | - | 12.8 | 0.01 |
| Chlorophyll | - | - | 13.0 | 0.01 | - | - | 13.5 | 0.01 |
| Anthocyanins ( Abs g.DM<sup>-1</sup>) | 0.0676 | 28.7 | - | <0.01 | 0.024 | 12.5 | - | 0.01 |

\*MS= Mean square, F= F-value, H=Kruskal-Wallis H-test, P= P-value

**Fig 5.**
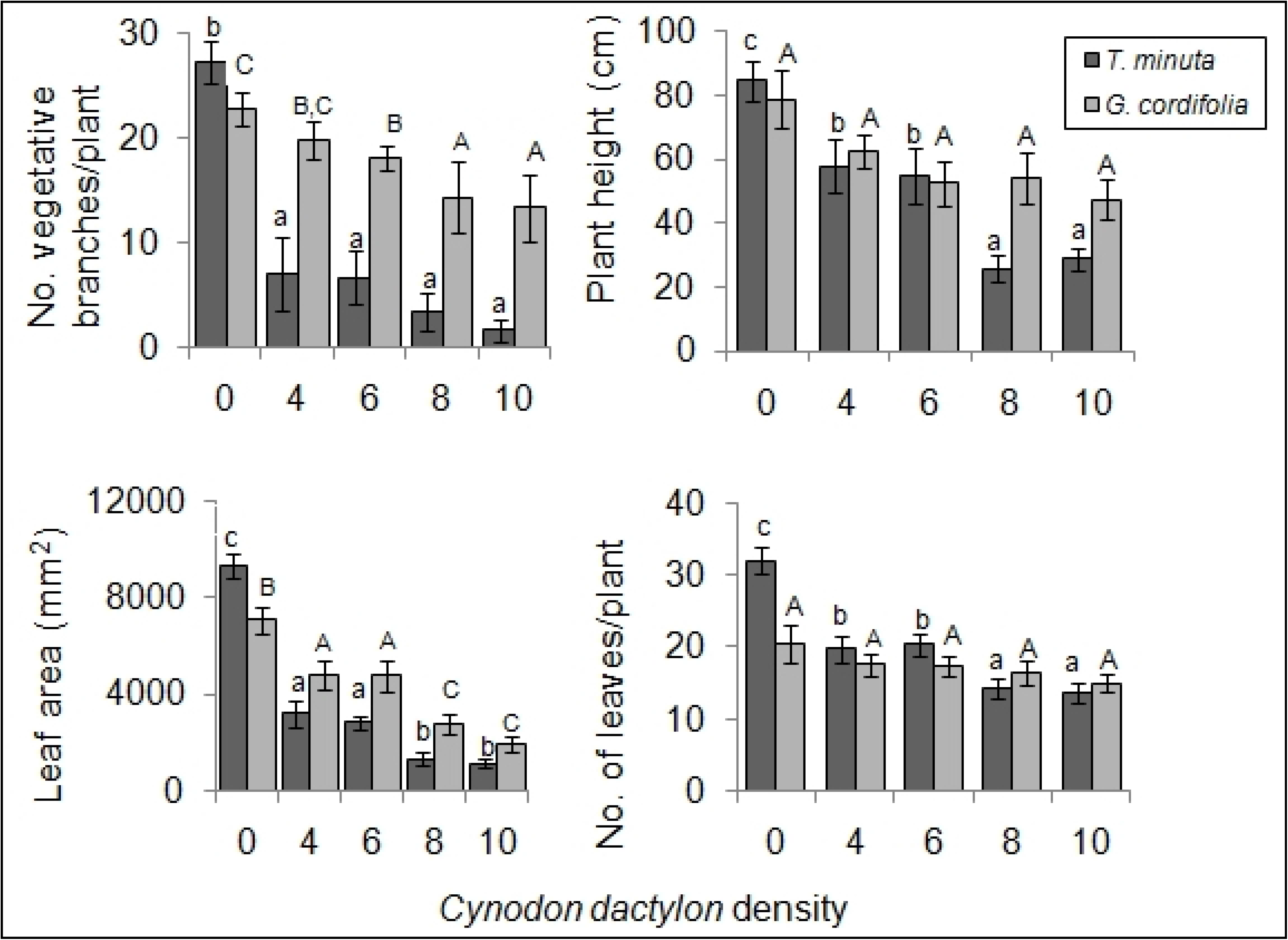
*Cynodon dactylon* varying density effects on the number of vegetative branches, plant height, leaf area and number of leaves of *T. minuta* and *G. cordifolia* in a screen house experiment. *Cynodon dactylon* densities ranged from 0 to 10 while the number of weeds per pot was two. Bars with dissimilar letters are significant by Fisher LSD at *p* = 0.05

**Fig 6.**
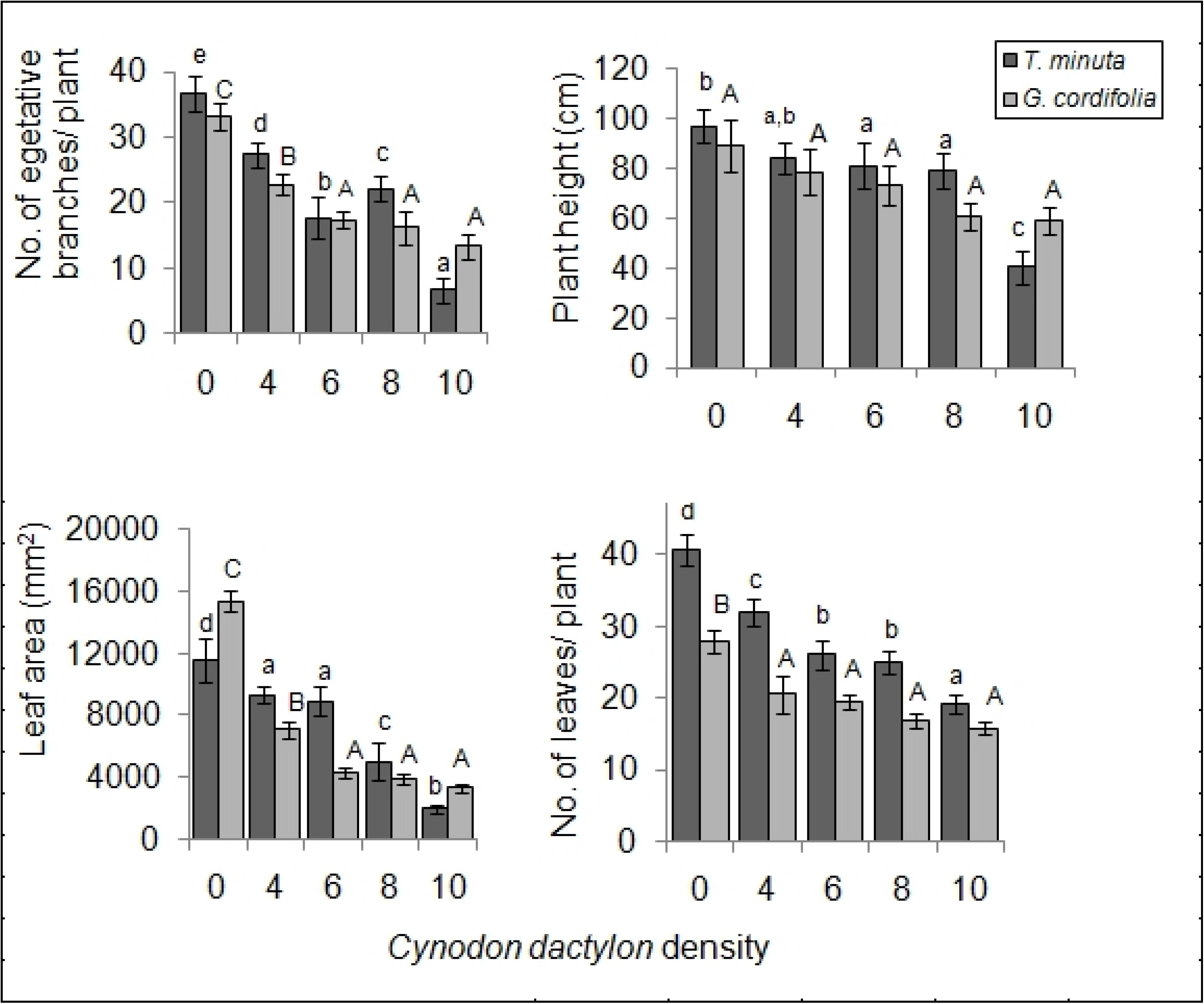
*Cynodon dactylon* varying density effects on the number of vegetative branches, plant height, leaf area and number of leaves of *T. minuta* and *G. cordifolia* in field plot experiment. *Cynodon dactylon* densities ranged from 0 to 10 while the number of weeds per plot was two. Bars with dissimilar letters are significant by Fisher LSD at *p* = 0.05

The mean number of leaves and plant height differed significantly in *T. minuta* (*p* < 0.05), being half as many and shorter in pots/plots with ≥ 8 *C. dactylon* per pot/plot as those in control treatment but no difference was observed for *G. cordifolia* (*p* > 0.05) (Figs 5 and 6).

Mean shoot diameter and shoot biomass differed significantly across the five *C. dactylon* densities in both *T. minuta* and *G. cordifolia* (*p* < 0.05). *Tagetes minuta* and *G. cordifolia* in pots/plots with *C. dactylon* density ≥ 8 per pot or plot had had half the diameter and were twice as light as *T. minuta* and *G. cordifolia* contained in control pots/plots. Mean root biomass and root length differed significantly only in *T. minuta* (*p* < 0.05) but not in *G. cordifolia* (*p* > 0.05) (Figs 7 and 8). *Tagetes minuta* in pots/plots with *C. dactylon* density ≥ 8 per pot or plot had roots with over four times lighter weight and half the length of roots of *T. minuta* in control pots/plots (Tables 1 and 2).

**Fig 7.**
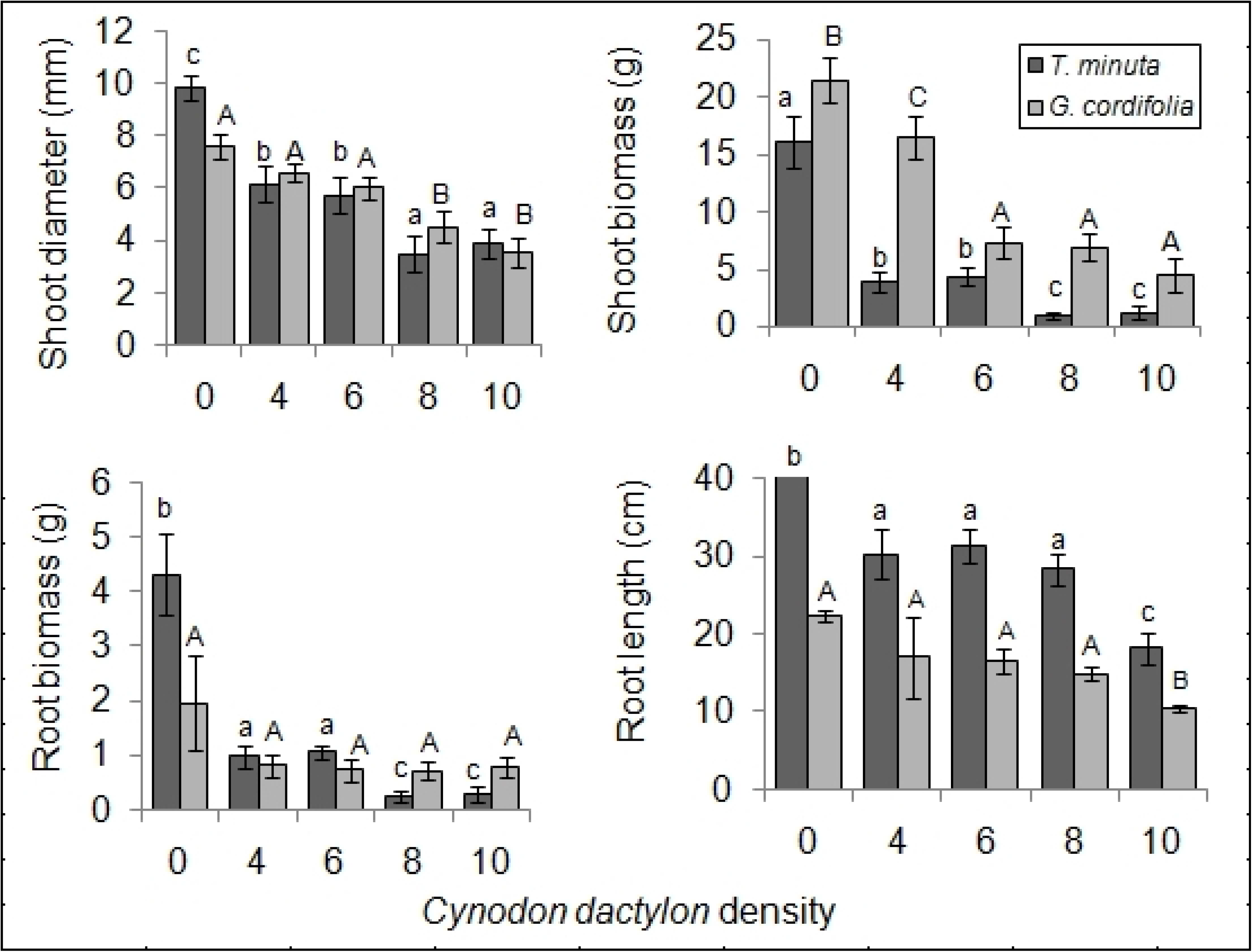
*Cynodon dactylon* varying density effects on mean shoot diameter, shoot biomass, root biomass, and root length of *T. minuta* and *G. cordifolia* intercrops in a screen house experiment. *Cynodon dactylon* densities ranged from 0 to 10 while the number of weeds per pot was two. Bars with dissimilar letters are significant by Fisher LSD at *p* = 0.05

**Fig 8.**
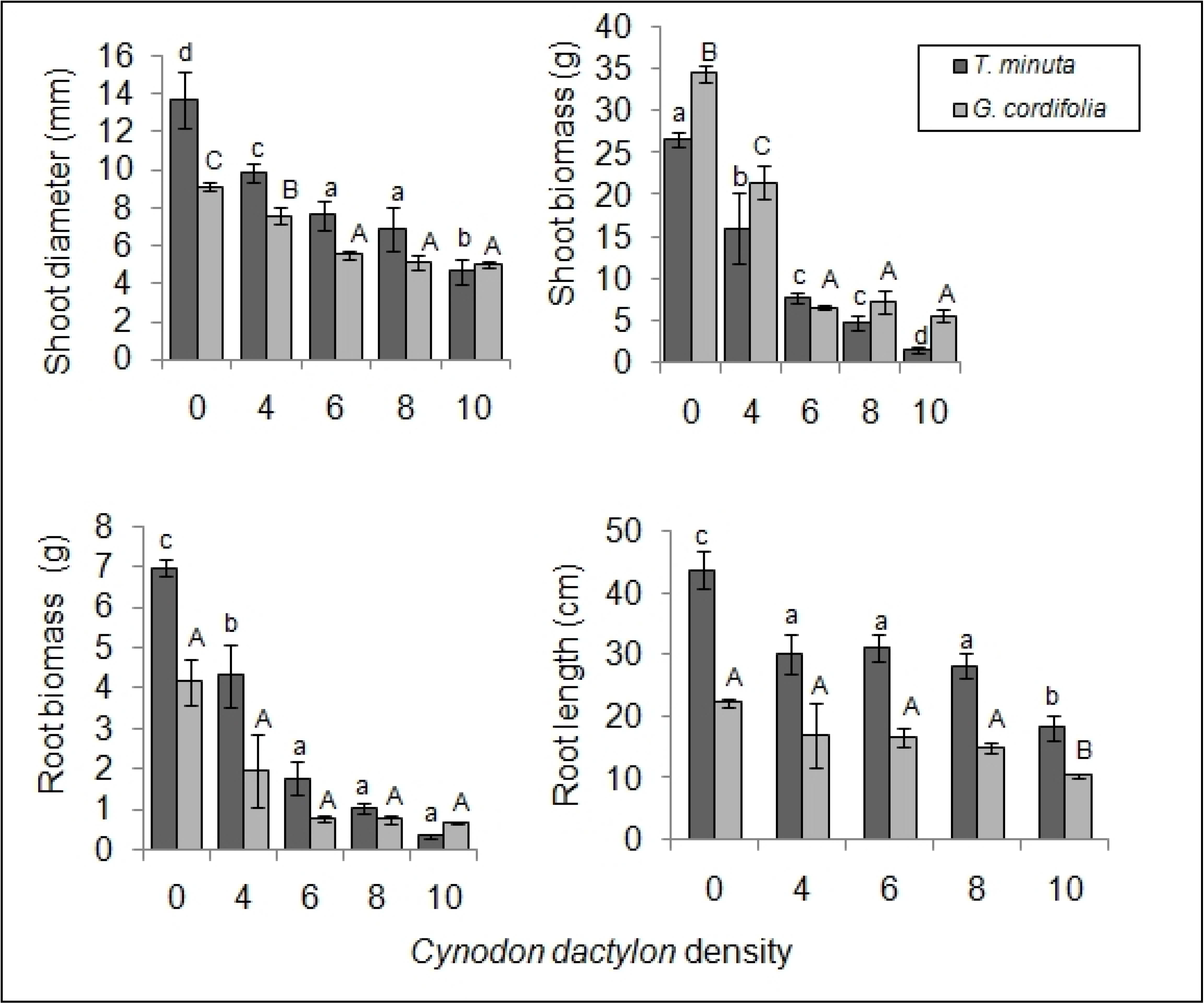
*Cynodon dactylon* varying density effects on mean shoot diameter, shoot biomass, root biomass, and root length of *T. minuta* and *G. cordifolia* intercrops in field plot experiment. *Cynodon dactylon* densities ranged from 0 to 10 while the number of weeds per plot was two. Bars with dissimilar letters are significant by Fisher LSD at *p* = 0.05

#### 3.1.3. Leaf pigmentations

In both *T. minuta* and *G. cordifolia*, total leaf chlorophyll content differed significantly across the five *C. dactylon* treatments that were planted separately (*p* < 0.05) (Fig 9; Tables 1 and 2). *Tagetes minuta* and *G. cordifolia* in control pots/plots had three times total leaf chlorophyll than *T. minuta* and *G. cordifolia* contained in pots/plots with *C. dactylon* density ≥ 8 per pot or plot.

**Fig 9.**
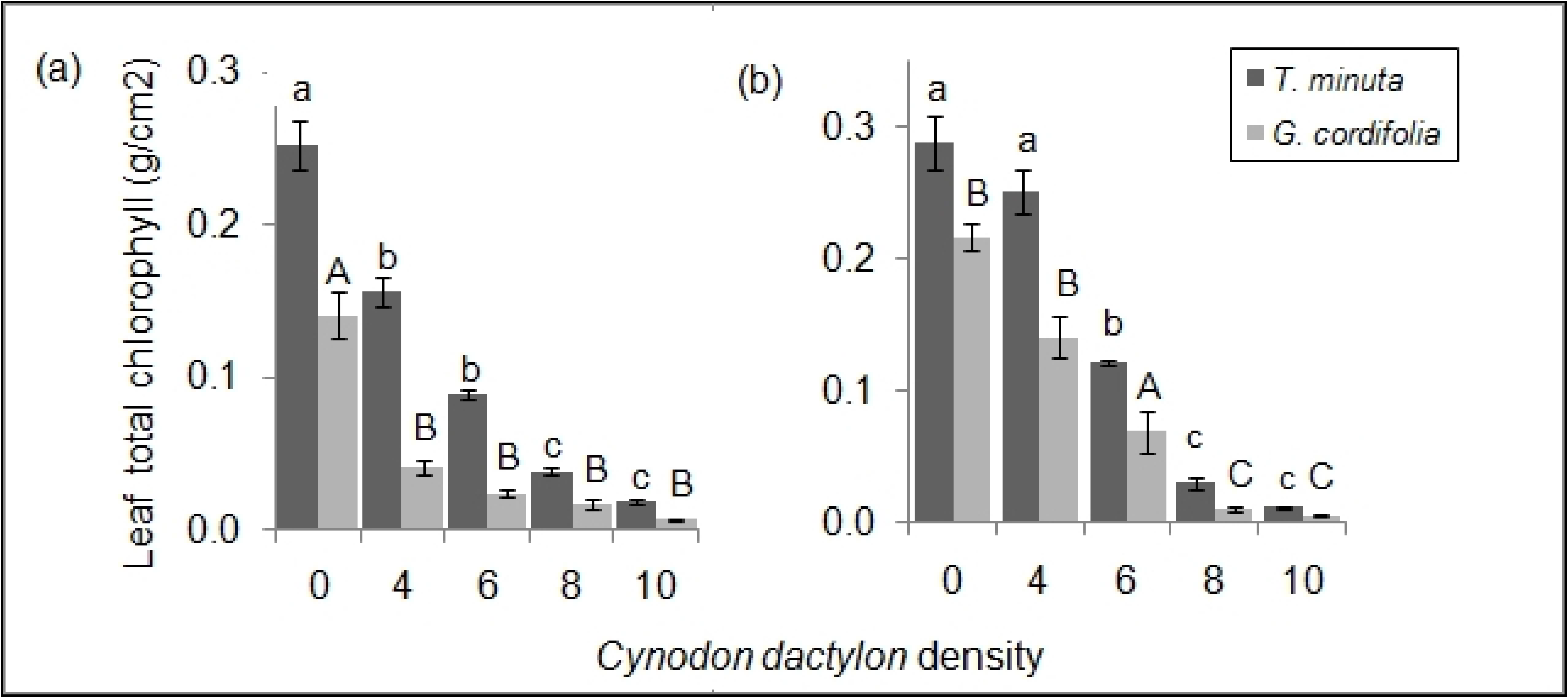
Mean (±SE) total leaf chlorophyll content of *T. minuta* and *G. cordifolia* planted with various *C. dactylon* densities (a) in screen house and (b) field plot experiments. *Cynodon dactylon* densities ranged from 0 to 10 while the number of weeds per pot/plot was two

Leaf anthocyanin concentrations differed significantly across the five *C. dactylon* treatments in both *T. minuta* and *G. cordifolia* (*p* < 0.05) (Fig 10; Tables 1 and 2). *Tagetes minuta* and *G. cordifolia* contained in pots/plots with *C. dactylon* density ≥ 8 per pot or plot had twice the total leaf anthocyanin than *T. minuta* and *G. cordifolia* in control pots/plots.

**Fig 10.**
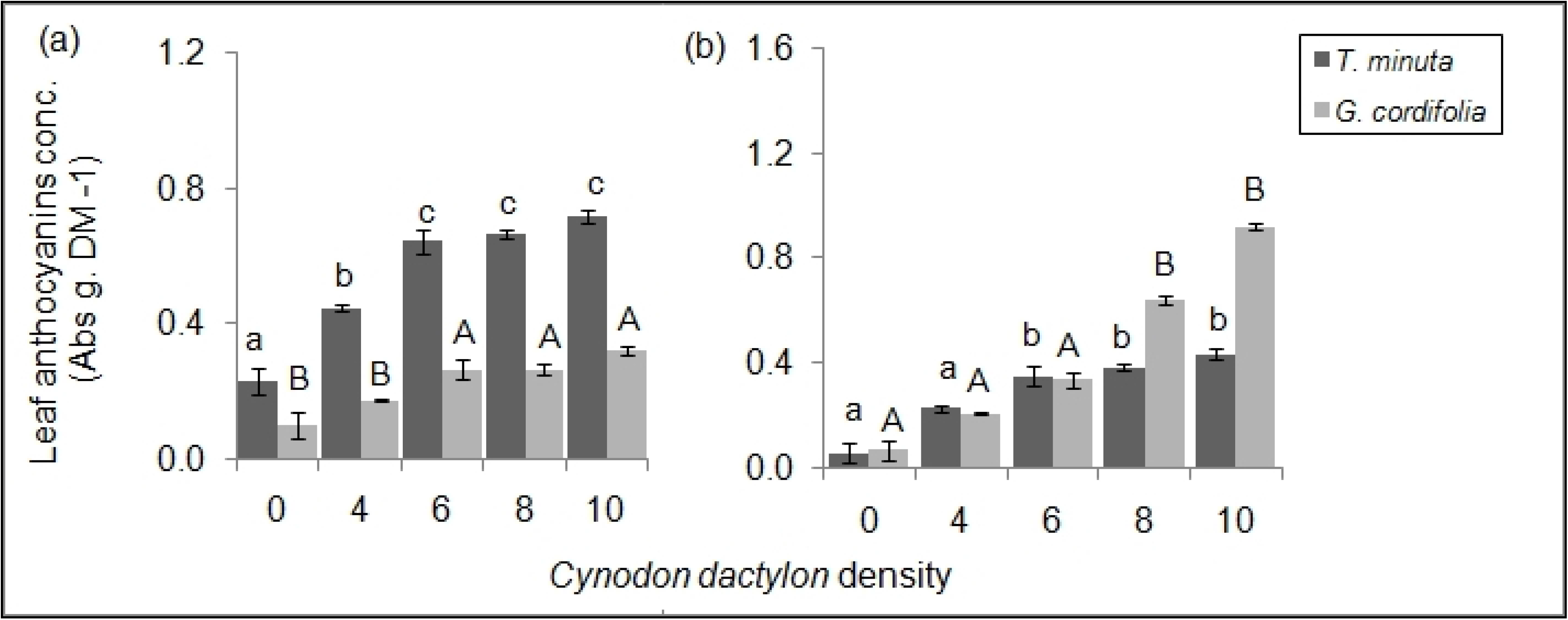
Mean (±SE) total leaf anthocyanins of *T. minuta* and *G. cordifolia* planted with various *C. dactylon* densities (a) in screen house and (b) field plot experiments. *Cynodon dactylon* densities ranged from 0 to 10 while the number of weeds per pot/plot was two

## 4.0. Discussion

### 4.1. Cynodon dactylon density dependent competitive effects on T. minuta and G. cordifolia growth parameters

The experiments showed that increasing densities of *C. dactylon* affects *T. minuta* and *G. cordifolia* negatively. While increasing densities of *C. dactylon* decreased plant height, shoot diameter, shoot biomass, leaf area, root biomass, the number of vegetative branches, leaves and root length of *T. minuta*, only shoot diameter, shoot biomass, the number of vegetative branches and leaf area were affected in *G. cordifolia*. These growth parameters are crucial to the growth and overall fitness of a plant [pers. obs.]. Plant height, leaf area and number of leaves for instance have ensured plants to intercept up to a recommended 95% of the incoming solar radiation for photosynthesis [35,36]. While shoot diameter and biomass aid in overcoming stresses (stress tolerance) such as trampling by animals and wind destruction [37], the number of panicles per plant determines the amount of seeds deposited in the soil which is crucial for invasion success of most weeds [38]. We observed a significant reduction in the number of *T. minuta* & *G. cordifolia* panicles per increasing density of *C. dactylon* which signifies that *C. dactylon* potentially reduce *T. minuta* and *G. cordifolia* seeds to be deposited in invaded areas. Therefore, the observed evidence suggest that; reduction in of panicles is likely to reduce seed production by weeds, thus allowing for application of this technique in the management of weeds in affected areas [38,39]. The greater negative effects of *C. dactylon* on *T. minuta* versus *G. cordifolia* may be due to *T. minuta*’s shorter and lighter roots compared to those of G. cordifolia [pers. obs]. Possibly, *G. cordifolia*’s greater root weight and length render this species less prone to suppression by competition compared to *T. minuta*. As predicted, the negative competitive effects were more pronounced with increasing densities of *C. dactylon*, a plant that has been reported as a very strong competitor to most crops [13] likely due to increased competition for available nutrients and space, in which *C. dactylon* out-competed the two invasive weeds. Competitiveness of *C. dactylon* has been associated with its stoloniferous nature and an ability to develop deep roots [9] that easily escape the effects competition. While monocultures from invasive plants have been reported to not only suppress other native species [40] but also bad for soil health [41]; intercrops were shown to be mostly of facilitative nature, especially in maize-legume combinations [42,43]. In contrast, in our study particularly the *C. dactylon* density dependent competition resulting from inter-planting *C. dactylon* / *T. minuta* and or *C. dactylon* /*G. cordifolia* can potentially reduce invasiveness of the two weeds as weed’ growth parameters that are crucial for plant fitness were significantly reduced.

### 4.2. Cynodon dactylon density dependent competitive effects on T. minuta and G. cordifolia leaf pigmentations

As *C. dactylon* density increased, leaf chlorophyll content dropped in both *T. minuta* and G. *cordifolia*. This could be due to the weed species’ reduced access to water, nutrients and space. For example, as Nitrogen becomes less available to a particular plant, its chlorophyll production is reduced [44,45] and, consecutively, its leaf chlorophyll content. The results of this study imply that an increase in *C. dactylon* density has a potential of exerting enough stress to affect the two weeds’ chlorophyll productivity. Leaf chlorophyll content has been linked to plant health status [31] as it is associated with energy production and, hence, important for other metabolic activities [31]. Plants with reduced chlorophyll amount and, thereby, reduced photosynthetic capacity [45] also possess flowers with accelerated abscission [46], which reduces chances of dispersal by pollinators. Reduced dispersal of the two weeds will reduce the chance for weed’s monoculture formation, which has been proven to be devastating in an invaded ecosystem [40,47]. Increasing density of *C. dactylon* in *T. minuta* and or *G. cordifolia* invaded areas therefore, can potentially be used as an environmentally friendly invasive species management approach.

We observed an increasing anthocyanin concentration in *T. minuta* and *G. cordifolia* with increasing numbers of *C. dactylon*. Anthocyanins, which are a small group of pigments within flavonoids, form red-blue coloration in most plants [48]. The increase of anthocyanin levels in plant leaves under increasing *C. dactylon* densities in this study can be linked to the increasing level of competition, specifically for nutrients and space due to increasing density of *C. dactylon*. This is in line with [49] who argued that anthocyanin induction and / or accumulation in a plant tissue can be associated with nitrogen and / or phosphorus deficiency. *Cynodon dactylon* competitive effects as a strategy for suppressing *T. minuta* and *G. cordifolia* [50] could be another possible cause of increased anthocyanins in leaves of both *T. minuta* and *G. cordifolia* exposed to increasing density of *C. dactylon*. Generally, the presence of these pigmentations in leaves is normally associated with stressors [48]. In this study, the stressor that have possibly induced increased anthocyanins in leaves of both *T. minuta* and *G. cordifolia* can be associated with competitive effects of *C. dactylon* for the available nutrients, space and water. It is a known fact that the rate of photosynthesis is directly proportional to plant’s chlorophyll content [51] intercepting solar radiation. Anthocyanin pigments reduce a plant’s chlorophyll content, thereby negatively affecting photosynthesis. Therefore, we claim that treating the two weeds with increasing densities of *C. dactylon* can be used to biologically manage the two invasive weeds efficiently.

## 5.0. Conclusions

In this study, shorter plant height, smaller shoot diameter, smaller leaf area and lower shoot biomass of *T. minuta* and *G. cordifolia* under higher *C. dactylon* densities reduces both *T. minuta* and *G. cordifolia* fitness. Moreover, reduced leaf total chlorophyll and increased anthocyanin levels in leaves affects the photosynthetic ability of both invasives *T. minuta* and *G. cordifolia*. The net effect, therefore, is the development of weaker *T. minuta* and *G. cordifolia* plants that are easily affected by other stressors such as animal trampling and, thus, can be managed accordingly. Our screen house and field experiments determined that the critical density of *C. dactylon* to suppress the two invasive species studies is ≥ 8 plants/m^2^. Our findings show an alternative way to suppress weeds, particularly in rangelands and protected areas.

## Acknowledgements

Much appreciation is expressed to all laboratory technicians of the Nelson Mandela African Institution of Science and Technology for facilitating the conduction of the experiments.

